# Mild respiratory SARS-CoV-2 infection can cause multi-lineage cellular dysregulation and myelin loss in the brain

**DOI:** 10.1101/2022.01.07.475453

**Authors:** Anthony Fernández-Castañeda, Peiwen Lu, Anna C. Geraghty, Eric Song, Myoung-Hwa Lee, Jamie Wood, Belgin Yalçın, Kathryn R. Taylor, Selena Dutton, Lehi Acosta-Alvarez, Lijun Ni, Daniel Contreras-Esquivel, Jeff R. Gehlhausen, Jon Klein, Carolina Lucas, Tianyang Mao, Julio Silva, Mario A. Peña-Hernández, Alexandra Tabachnikova, Takehiro Takahashi, Laura Tabacof, Jenna Tosto-Mancuso, Erica Breyman, Amy Kontorovich, Dayna McCarthy, Martha Quezado, Marco Hefti, Daniel Perl, Rebecca Folkerth, David Putrino, Avi Nath, Akiko Iwasaki, Michelle Monje

## Abstract

Survivors of Severe Acute Respiratory Syndrome Coronavirus-2 (SARS-CoV-2) infection frequently experience lingering neurological symptoms, including impairment in attention, concentration, speed of information processing and memory. This long-COVID cognitive syndrome shares many features with the syndrome of cancer therapy-related cognitive impairment (CRCI). Neuroinflammation, particularly microglial reactivity and consequent dysregulation of hippocampal neurogenesis and oligodendrocyte lineage cells, is central to CRCI. We hypothesized that similar cellular mechanisms may contribute to the persistent neurological symptoms associated with even mild SARS-CoV-2 respiratory infection. Here, we explored neuroinflammation caused by mild respiratory SARS-CoV-2 infection – without neuroinvasion - and effects on hippocampal neurogenesis and the oligodendroglial lineage. Using a mouse model of mild respiratory SARS-CoV-2 infection induced by intranasal SARS-CoV-2 delivery, we found white matter-selective microglial reactivity, a pattern observed in CRCI. Human brain tissue from 9 individuals with COVID-19 or SARS-CoV-2 infection exhibits the same pattern of prominent white matter-selective microglial reactivity. In mice, pro-inflammatory CSF cytokines/chemokines were elevated for at least 7-weeks post-infection; among the chemokines demonstrating persistent elevation is CCL11, which is associated with impairments in neurogenesis and cognitive function. Humans experiencing long-COVID with cognitive symptoms (48 subjects) similarly demonstrate elevated CCL11 levels compared to those with long-COVID who lack cognitive symptoms (15 subjects). Impaired hippocampal neurogenesis, decreased oligodendrocytes and myelin loss in subcortical white matter were evident at 1 week, and persisted until at least 7 weeks, following mild respiratory SARS-CoV-2 infection in mice. Taken together, the findings presented here illustrate striking similarities between neuropathophysiology after cancer therapy and after SARS-CoV-2 infection, and elucidate cellular deficits that may contribute to lasting neurological symptoms following even mild SARS-CoV-2 infection.

## Introduction

The COVID-19 pandemic caused by severe acute respiratory syndrome coronavirus-2 (SARS-CoV-2) has to date resulted in over 285 million documented COVID-19 cases worldwide. Neurological symptoms are emerging as relatively common sequelae of SARS-CoV-2 infection, with persistent cognitive impairment affecting approximately one in four COVID-19 survivors (Nasserie et al., 2021). While more common in individuals who had experienced severe COVID requiring hospitalization, even those with mild symptoms in the acute phase may experience lasting cognitive dysfunction (Becker et al., 2021; Nasserie et al., 2021). Colloquially known as “COVID-fog”, this syndrome of COVID-associated cognitive impairment is characterized by impaired attention, concentration, speed of information processing, memory, and executive function (Becker et al., 2021; Nasserie et al., 2021). Together with increased rates of anxiety, depression, disordered sleep and fatigue, this syndrome of cognitive impairment contributes substantially to the morbidity of “long-COVID” and in many cases prevents people from returning to their previous level of occupational function (Davis et al., 2021; Tabacof et al., 2022). Given the scale of SARS-CoV-2 infection, this syndrome of persistent cognitive impairment represents a major public health crisis (Nath 2020).

The syndrome of cognitive symptoms that COVID survivors frequently experience closely resembles the syndrome of cancer therapy-related cognitive impairment (CRCI), commonly known as “chemo-brain”. Neuroinflammation is central to the pathophysiology of cancer therapy-related cognitive impairment (for review, see (Gibson and Monje, 2021)), raising the possibility of shared pathophysiological mechanisms. Microglia, brain-resident macrophages, become persistently reactive following exposure to certain systemic chemotherapy drugs, after cranial radiation, or after systemic inflammatory challenge with low-dose lipopolysaccharide (Geraghty et al., 2019; Gibson et al., 2019; Monje et al., 2002; Monje et al., 2003; Monje et al., 2007). A distinct subpopulation of microglia that reside in white matter (Hammond et al., 2019) are selectively activated by systemic insults such as exposure to the chemotherapy drug methotrexate (Gibson et al., 2019). Reactive microglia impair mechanisms of cellular homeostasis and plasticity such as the ongoing generation of myelin-forming oligodendrocytes (Gibson et al., 2019), myelin plasticity (Geraghty et *al*., 2019) and new neuron generation in the hippocampus (Monje et al., 2002; Monje et al., 2003; Monje et al., 2007). Local microglial cytokine secretion contributes to at least part of this dysregulation (Monje et al., 2003). Elevated circulating cytokine/chemokine levels, particularly CCL11, can also limit neurogenesis and contribute to cognitive impairment (Villeda et al., 2011). In addition to these direct effects of inflammatory mediators on cellular plasticity in the brain, microglia also induce neurotoxic astrocyte reactivity through cytokine signaling (Liddelow et al., 2017). Astrocytes can assume a range of reactive states (Hasel et al., 2021) that can induce further pathophysiology, with certain states of reactive astrocytes inducing oligodendrocyte and neuronal cell death (Liddelow et al., 2017) through secretion of saturated lipids contained in lipoprotein particles (Guttenplan et al., 2021). This complex cellular dysregulation is thought to contribute importantly to cognitive impairment, and anti-inflammatory strategies correct such multicellular dysregulation and rescue cognition in disease states such as occurs after exposure to neurotoxic cancer therapies (Geraghty et al., 2019; Gibson et al., 2019; Monje et al., 2003) and in aging (Villeda et al., 2011).

While systemic and severe COVID can cause multi-organ disease and numerous potential mechanisms affecting the nervous system (Lee et al., 2021; Nath and Smith, 2021; Remsik et al., 2021), even mild COVID could result in an a detrimental neuroinflammatory response, including reactivity of these exquisitely sensitive white matter microglia. Given this context, we hypothesized that the inflammatory response to even mild COVID-19 may induce elevation in neurotoxic cytokines/chemokines, a pattern of white matter microglial reactivity, and consequent dysregulation of myelin-forming oligodendrocytes and hippocampal neural precursor cells.

### Mild respiratory SARS-CoV-2 infection causes prominent neuroinflammation

To test the effects of mild COVID-19, we used a mouse model of mild SARS-CoV-2 infection limited to the respiratory system (Israelow et al., 2020; Song et al., 2021). SARS-CoV-2 infection requires expression of human ACE2. In this model, human ACE2 is delivered via AAV vector to the trachea and lungs. Two weeks following intra-tracheal ACE2-AAV delivery, SARS-CoV-2 is delivered intranasally (Figure 1A). Control mice received intra-tracheal AAV expressing human ACE2, but only mock infection intranasally. Mice exhibit no weight loss (Figure 1B) or observable sickness behavior; the challenges of working with a biosafetly level-3 mouse model limited the feasibility of further behavioral assessments. No virus was detected in brain (Figure 1C), unlike in the case of a neuroinvasive model of SARS-CoV-2 infection (Supplementary Figure 1). As expected, SARS-CoV-2 was present in the lungs of infected mice (Supplementary Figure 2). Despite the lack of evident symptoms/signs of illness in this respiratory infection model, we found prominently elevated cytokine profiles in serum and in cerebrospinal fluid (CSF) at 7-day and 7- week timepoints following respiratory infection, in comparison to control mice (Figure 1D-G). At 7-days post-infection, elevated CSF cytokines and chemokines include CXCL10, IL6, IFN-g, CCL7, CCL2, CCL11 and BAFF, among others (Figure 1F). Of these, CXCL10, CCL7 and CCL11 remain elevated in CSF at 7-weeks post-infection (Figure 1G). These results indicate that respiratory infection with SARS-CoV-2 results in profound changes in cytokines within the CSF.

**Figure 1.**
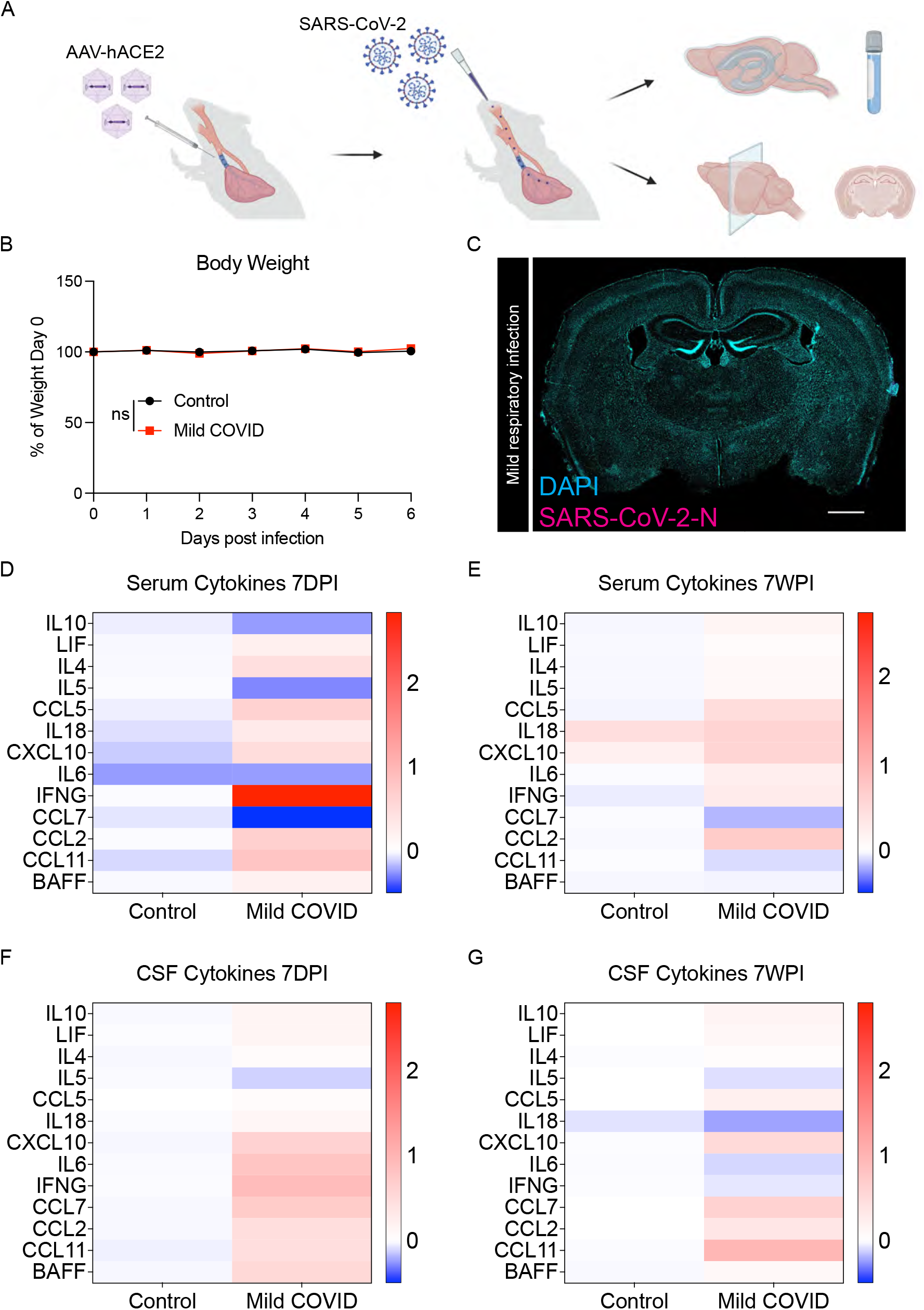
Mouse model of mild respiratory SARS-CoV-2 infection. (A) Schematic of experimental paradigm for respiratory system-restricted SARS-CoV-2 infection in mice and experimental workflow, illustrating intratracheal AAV-hACE2 delivery, SARS-CoV- 2 intranasal infection, CSF collection, and tissue processing for microscopy analysis. (B) Body weight (% of day 0 weight) of control and mild COVID mice. Data shown as mean +/- SEM; n=24 mice per group; ns p>0.05 by 2-way ANOVA with multiple comparisons. (C) Confocal micrograph of SARS-CoV-2 nucleocapsid protein (SARS-CoV-2-N) 7-days post-infection (7DPI). SARS-CoV-2-N, magenta; DAPI, cyan. Scale bar = 1mm. (D and E) Cytokine analyses of serum in control and mild COVID mice 7-days post-infection (D) and 7-weeks post-infection (7WPI) (E). Data shown as mean log_2_(fold change mean fluorescence intensity) normalized to control group; n=7 CD1 mice per group. (F and G) Cytokine analysis of CSF in control and mild COVID mice 7-days post-infection (F) and 7-weeks post-infection (G). Data shown as mean log_2_(fold change mean fluorescence intensity) normalized to control group; n=7 CD1 mice per group.

We next examined microglial reactivity in the mild COVID mouse model compared to control mice. As hypothesized, we found increased microglial reactivity in subcortical white matter but not in cortical gray matter, as assessed by IBA1 and CD68 co-positivity (Figure 2). This pattern of white matter-specific microglial reactivity was evident in two different mouse strains (CD1 and BALB/c) at 7-days post-infection (Figure 2A-C) and persisted to 7-weeks post-infection (Figure 2D-F). In contrast to the subcortical white matter, microglial reactivity was not observed in the cortex (Figure 2G-L).

**Figure 2.**
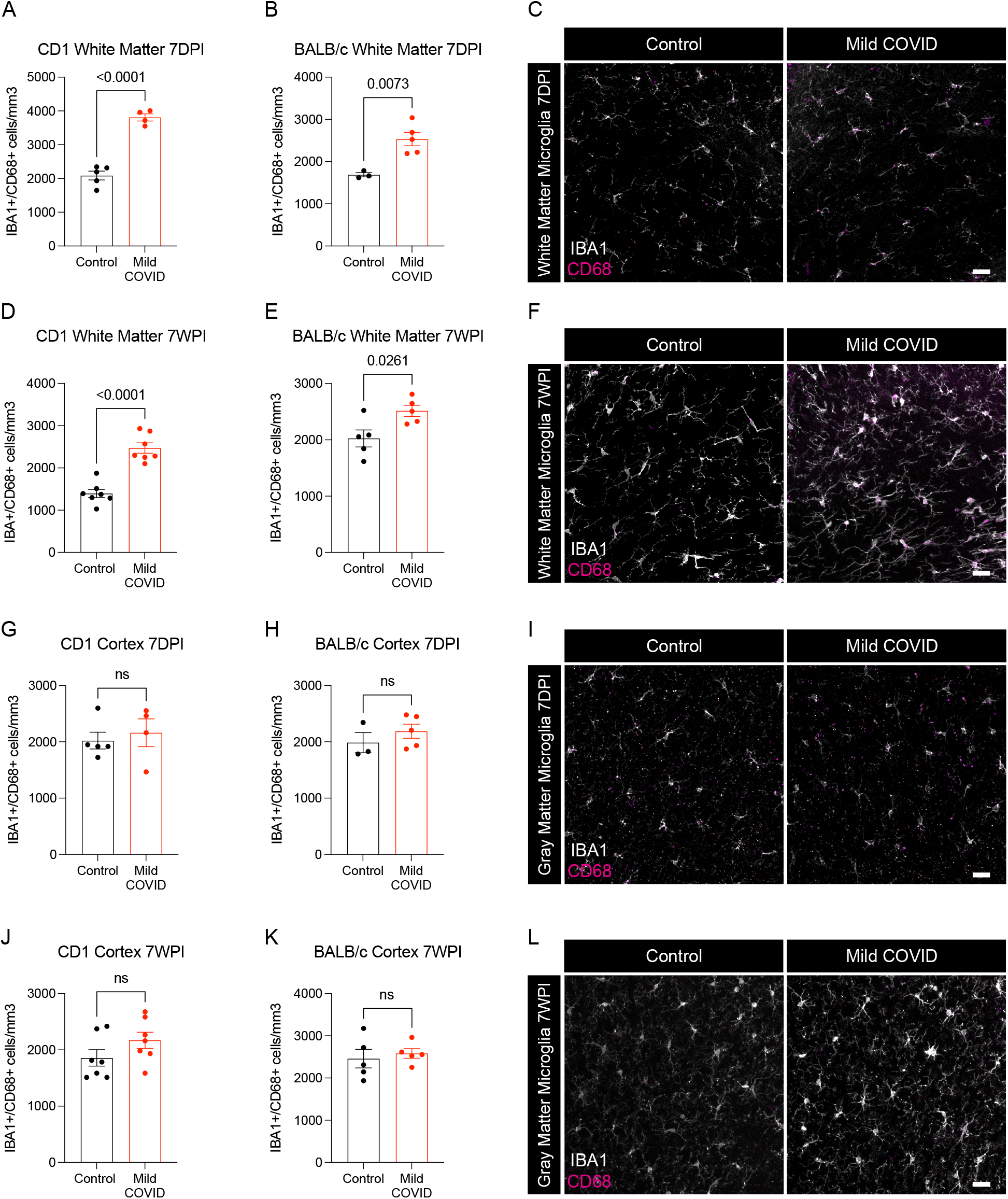
White matter microglial activation after mild respiratory SARS-CoV-2 infection. (A and B) Activated microglia (IBA1^+^ CD68^+^) quantification 7-days post-infection in the cingulum of the corpus callosum of CD1 (A) and BALB/c (B) mice. n=5 mice per CD1 control group; n=4 mice per CD1 mild COVID group; n=3 mice per BALB/c control group; n=5 mice per BALB/c mild COVID group. (C) Representative confocal micrographs of activated microglia (IBA1, white; CD68, magenta) in the cingulum of the corpus callosum of BALB/c mice 7-days post-infection. (D and E) Activated microglia (IBA1^+^ CD68^+^) quantification 7-weeks post-infection in the cingulum of the corpus callosum of CD1 (D) and BALB/c (E) mice. n=7 mice per CD1 control group; n=7 mice per CD1 mild COVID group; n=5 mice per BALB/c control group; n=5 mice per BALB/c mild COVID group. (F) Representative confocal micrographs of activated microglia (IBA1, white; CD68, magenta) in the cingulum of the corpus callosum of BALB/c mice 7-weeks post-infection. (G and H) Activated microglia (IBA1^+^ CD68^+^) quantification 7-days post-infection in the cortical gray matter of CD1 (G) and BALB/c (H) mice. n=5 mice per CD1 control group; n=4 mice per CD1 mild COVID group; n=3 mice per BALB/c control group; n=5 mice per BALB/c mild COVID group. (I) Representative confocal micrographs of activated microglia (IBA1, white; CD68, magenta) in the cortical gray matter of BALB/c mice 7-days post-infection. (J and K) Activated microglia quantification (IBA1^+^ CD68^+^) 7-weeks post-infection in the cortical gray matter of CD1 (J) and BALB/c (K) mice. n=7 mice per CD1 control group; n=7 mice per CD1 mild COVID group; n=5 mice per BALB/c control group; n=5 mice per BALB/c mild COVID group. (L) Representative confocal micrographs of activated microglia (IBA1, white; CD68, magenta) in the cortical gray matter of BALB/c mice 7-weeks post-infection. Data shown as mean +/- SEM; each dot represents an individual mouse; P values shown on figure panels; ns p>0.05 by two-tailed, unpaired t-test. Scale bars 50μm.

We had the opportunity to examine human cortex and subcortical white matter samples from a cohort of nine individuals (7 male, 2 female, age 24-73) found to be SARS-CoV-2-positive by nasal swab PCR at the time of death during the spring of 2020 (March – July 2020). Five of these subjects died suddenly and were found to have SAR-CoV-2 infection, and four subjects died within days to weeks after onset of symptoms; of these 9 subjects only one individual required ICU admission (Lee et al., 2021). The control group represents a cohort of 5 subjects without SARS-CoV-2 infection (5 male, age 43-70). Similar to the mouse model, humans with COVID-19, including mild COVID-19 or asymptomatic SARS-CoV-2 infection, exhibit robustly elevated CD68+ microglial reactivity in subcortical white matter compared to cortical grey matter and compared to individuals without SARS-CoV-2 infection (Figure 3A-B, Supplementary Figure 3).

**Figure 3.**
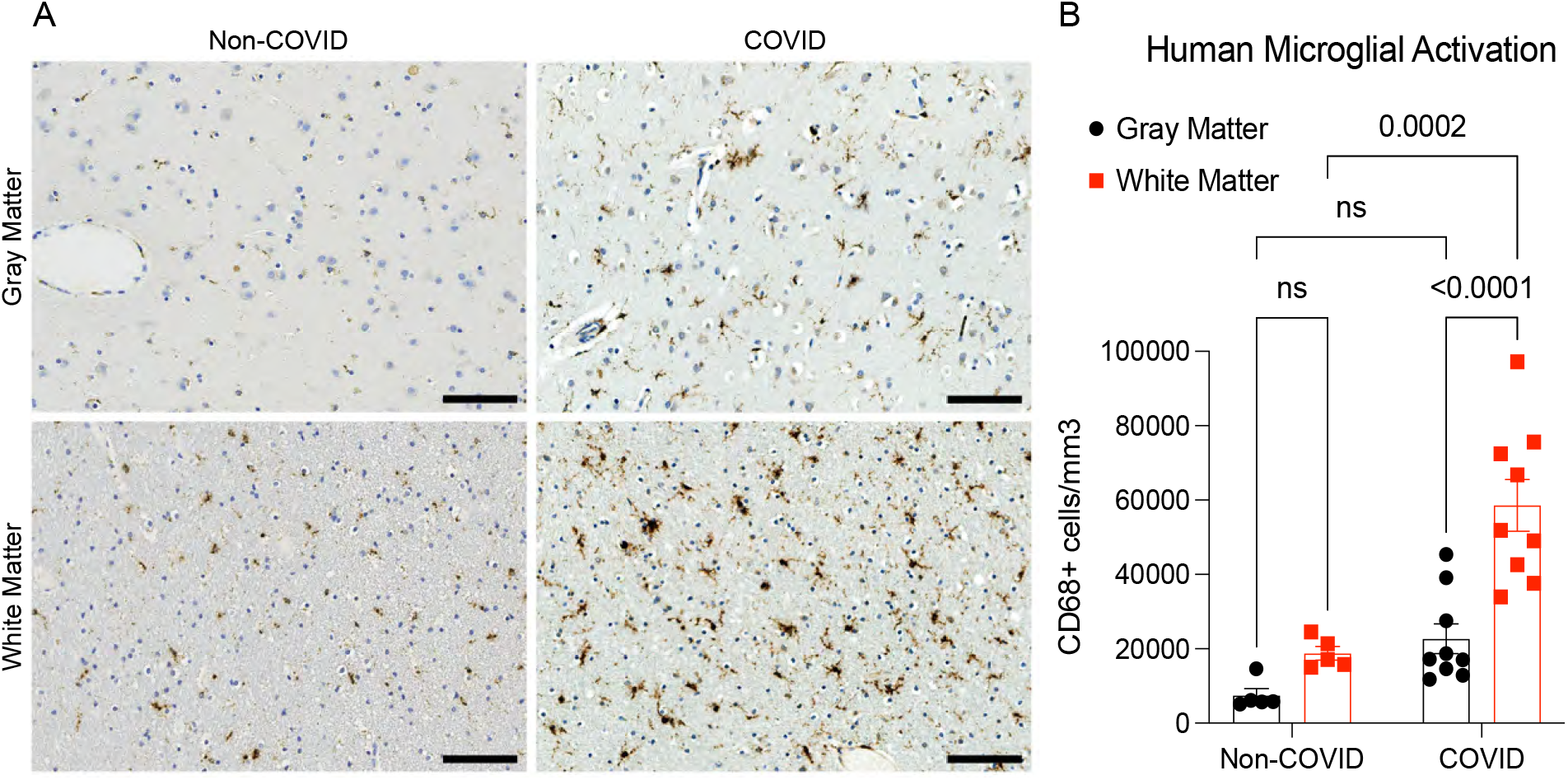
Microglial reactivity in human white matter after SARS-CoV-2 infection. (A) Representative micrographs of CD68 immunostaining (brown) in the gray (cerebral cortex) or subcortical white matter of human subjects with COVID or in non-COVID control subjects. (B) Activated microglia (CD68^+^ cells) quantification. n=5 for non-COVID control group, n=9 for COVID group. Data shown as mean +/- SEM; ns p>0.05 by 2-way ANOVA with multiple comparisons; each dot represents a human subject. Scale bars 100μm.

### Mild respiratory SARS-CoV-2 infection impairs hippocampal neurogenesis

Reactive microglia and other aspects of inflammatory responses to systemic illness (Monje et al., 2003) or aging (Villeda et al., 2011) can impair the generation of new neurons in the hippocampus, an ongoing mechanism of neural plasticity thought to support healthy memory function (Clelland et al., 2009; Zhang et al., 2008). Examining the mouse hippocampus following mild respiratory SARS-CoV-2 infection, we found robustly increased microglial reactivity in hippocampal white matter at 7-days post-infection (Figure 4A-B) that persists until at least 7-weeks post-infection (Figure 4C-D). Consistent with previous observations that reactive microglia can inhibit hippocampal neurogenesis (Monje et al., 2003), a stark decrease in new neuron generation was evident, as assessed by Doublecortin-positive cell quantification, at 7-days post-infection (Figure 4E-F) and persisted until at least 7-weeks post-infection (Figure 4G-H). Considering the number of newly generated hippocampal neurons as a function of the reactive microglial load, we found an inverse correlation between neurogenesis and reactive microglia in the hippocampus (Figure 4I).

**Figure 4.**
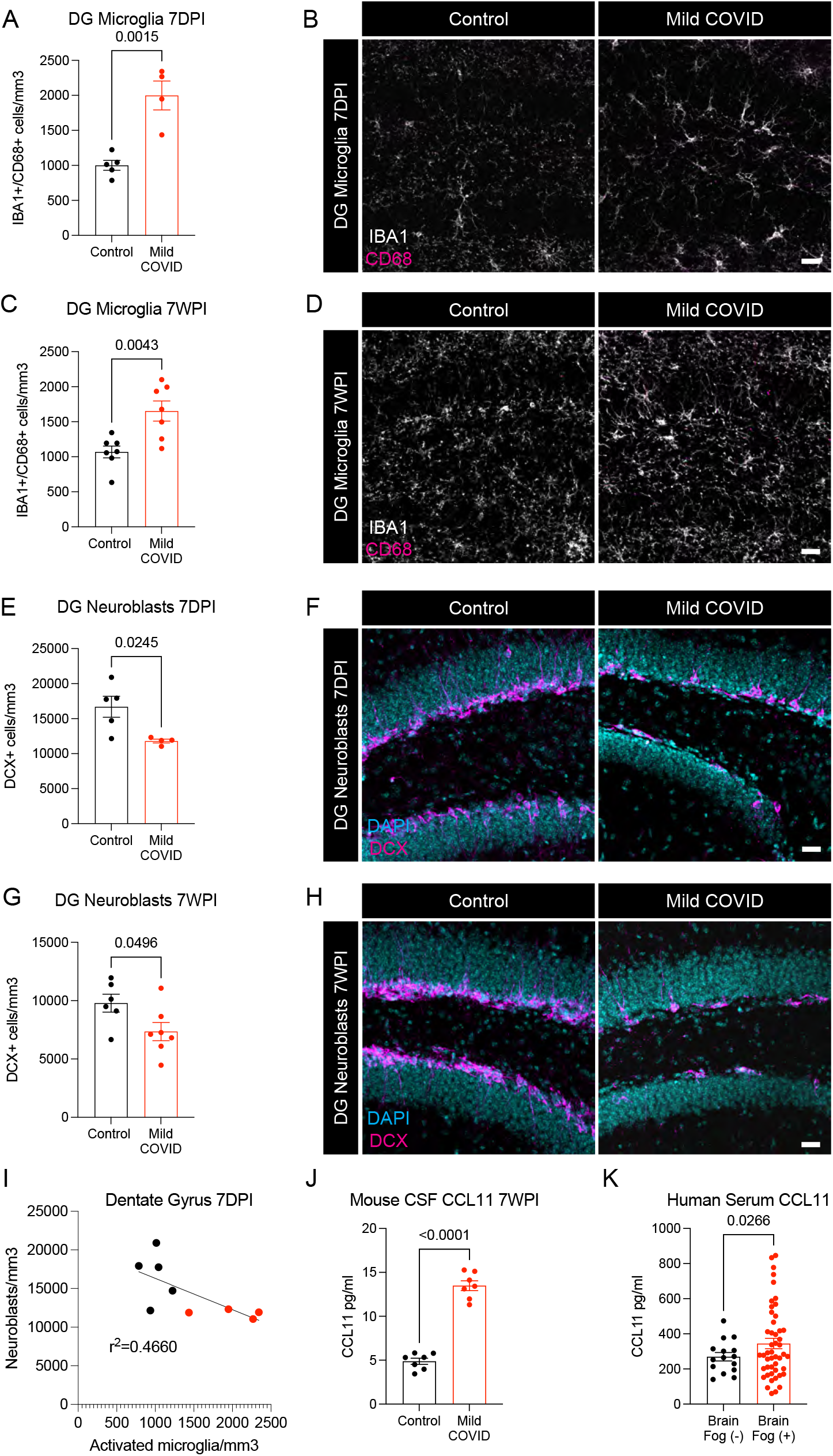
Decreased hippocampal neurogenesis after mild respiratory SARS-CoV-2 infection. (A) Activated microglia (IBA1^+^ CD68^+^) quantification 7-days post-infection in the dentate gyrus of CD1 mice. n=5 mice per control group; n=4 mice per mild COVID group. (B) Representative confocal micrographs of activated microglia (IBA1, white; CD68, magenta) in the dentate gyrus of CD1 mice 7-days post-infection. (C) Activated microglia (IBA1^+^ CD68^+^) quantification 7-weeks post-infection in the dentate gyrus of CD1 mice. n=7 mice per control group; n=7 mice per mild COVID group. (D) Representative confocal micrographs of activated microglia (IBA1, white; CD68, magenta) in the dentate gyrus of CD1 mice 7-weeks post-infection. (E) Neuroblast (DCX^+^) quantification 7-days post-infection in the dentate gyrus of CD1 mice. n=5 mice per control group; n=4 mice per mild COVID group. (F) Representative confocal micrographs of neuroblasts (DCX, magenta; DAPI, cyan) in the dentate gyrus of CD1 mice 7-days post-infection. (G) Neuroblast (DCX^+^) quantification 7-weeks post-infection in the dentate gyrus of CD1 mice. n=6 mice per control group; n=7 mice per mild COVID group. (H) Representative confocal micrographs of neuroblasts (DCX, magenta; DAPI, cyan) in the dentate gyrus of CD1 mice 7-weeks post-infection. (I) Correlation between neuroblasts (DCX^+^) and activated microglia (IBA1^+^ CD68^+^) in the dentate gyrus of CD1 mice 7-days post-infection. Line fitted with simple linear regression. n=5 mice per control group; n=4 mice per mild COVID group. (J) CCL11 levels in CSF of CD1 mice 7-weeks post-infection. n=7 mice per group. (K) Serum levels of CCL11 in people experiencing long-COVID with and without cognitive symptoms. n = 15 human subjects with long-COVID without cognitive symptoms (brain fog (-)), n = 48 human subjects with long-COVID with cognitive symptoms (brain fog (+)). Data shown as mean +/- SEM; each dot represents an individual mouse (A-J) or one human subject (K). A,C,E,G,J,K unpaired two-tailed t-test. P values shown on figure panels. Scale bars 50μm. DG: Dentate gyrus.

Inflammatory cytokines can directly inhibit hippocampal neurogenesis (Monje et al., 2003), including interleukin-6 (IL6) derived from reactive microglia (Monje et al., 2003) and the circulating cytokine CCL11 (also called eotaxin-1) (Villeda et al., 2011). CSF cytokine profiles at 7 days and 7 weeks following mild respiratory SARS-CoV-2 infection highlights elevated IL6 in CSF at 7 days, and persistently elevated CCL11 in CSF at 7 weeks. In the same mice, CCL11 was elevated at 7-days post-infection and – in contrast to persistent elevation in CSF – normalized in serum by 7-weeks post-infection (Figure 1D-G, Figure 4J and Supplementary Figure 4).

To further explore the relevance of elevated CCL11 levels to cognitive sequalae of COVID infection in humans, we next examined circulating CCL11 cytokine levels in the plasma of people suffering from long-COVID with and without cognitive symptoms. These individuals chiefly experienced a relatively mild course of SARS-CoV-2 infection, without hospitalization in more than 90% of subjects. We found elevated CCL11 levels in the plasma of people with long-COVID exhibiting cognitive deficits, or “brain fog” (n = 48 subjects, 16 male/32 female, mean age 46.1±14.6 years) compared to those with long-COVID lacking cognitive symptoms (n = 15 subjects, 4 male/11 female, mean age 46.7±14.1 years; Figure 4K).

### Mild respiratory SARS-CoV-2 infection impairs myelinating oligodendrocytes

We next explored the effects of SARS-CoV-2 mild respiratory infection on oligodendroglial lineage cells. Reactive microglia can alternatively promote (Miron et al., 2013) or impair (Gibson et al., 2019) oligodendrogenesis, depending on the precise microglial cell state. In disease states like CRCI, reactive microglia cause a dysregulation of the oligodendroglial lineage (Gibson et al., 2019) and loss of myelin plasticity (Geraghty et al., 2019). Examining oligodendroglial lineage cells in subcortical white matter (cingulum of the corpus callosum), we found maintenance of the oligodendrocyte precursor cell population marked by PDGFR-alpha at 7 days after mild respiratory SARS-CoV-2 infection (Figure 5A.) However, by 7 weeks following infection, a mild decrease (∼10%) in the number of oligodendrocyte precursor cells was evident (Figure 5B-C). Mature oligodendrocytes, assessed by ASPA and CC1 immunoreactivity, exhibited a greater magnitude of depletion (Figure 5D-F and Supplementary Figure 5), with loss of approximately one third of oligodendrocytes occurring by 7-days post-infection (Figure 5D). This depletion in mature oligodendrocytes persisted until at least 7-weeks post-infection (Figure 5E).

**Figure 5.**
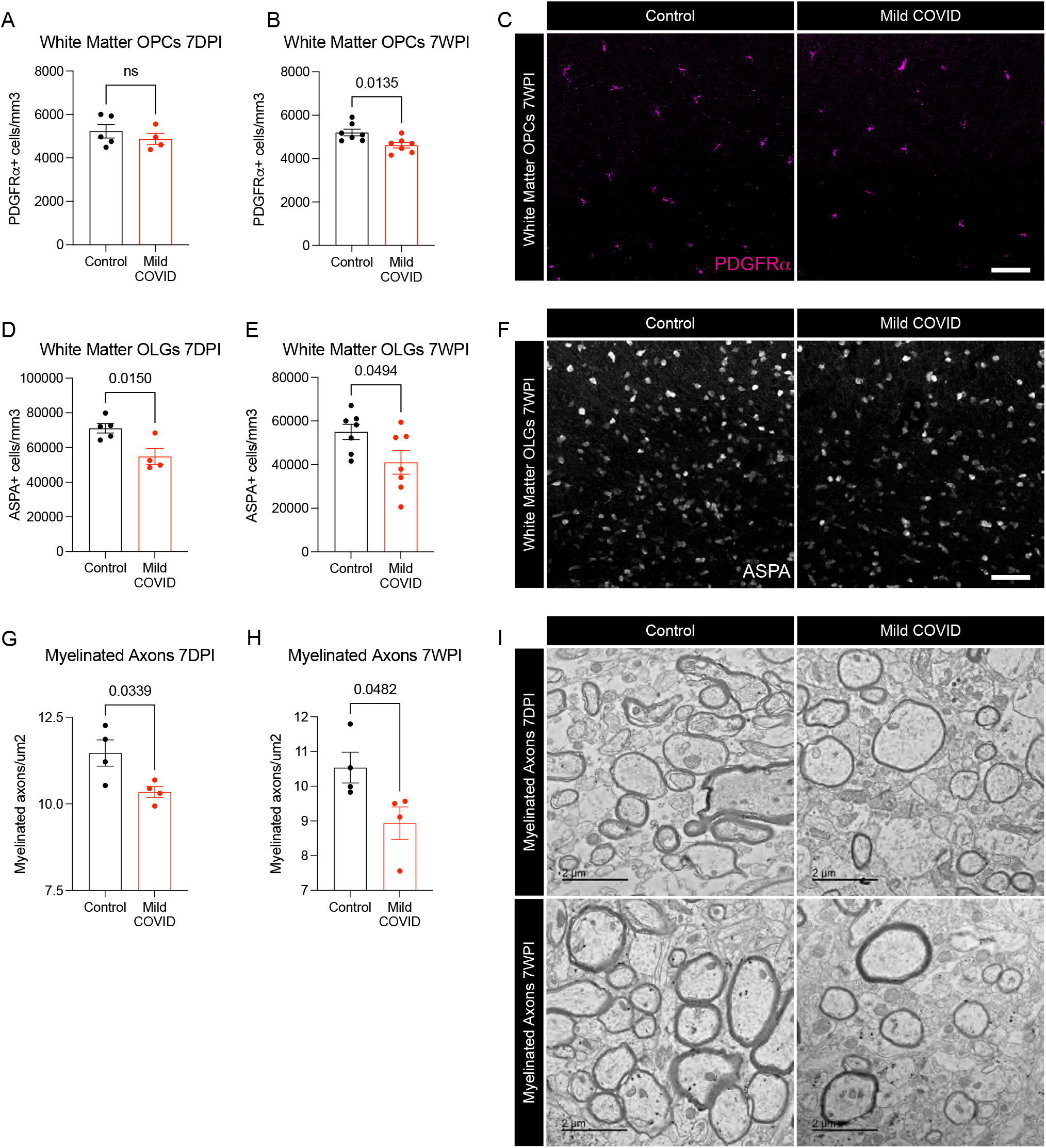
Oligodendrocyte and myelin loss after mild respiratory SARS-CoV-2 infection. (A and B) Oligodendrocyte precursor cell (PDGFRa^+^) quantification in the cingulum of the corpus callosum of CD1 mice 7-days (A) and 7-weeks (B) post-infection. n=5 mice per control group and n=4 mice per mild COVID group in (A). n=7 mice per group in (B). (C) Representative confocal micrographs of oligodendrocyte precursor cells (PDGFRa, magenta) in the cingulum of the corpus callosum of CD1 mice 7-weeks post-infection. Scale bar 50μm. (D and E) Oligodendrocyte (ASPA^+^) quantification in the cingulum of the corpus callosum of CD1 mice 7-days post-infection (D) and 7-weeks post-infection (E). n=5 mice per control group and n=4 mice per mild COVID group in (D). n=7 mice per group in (E). (F) Representative confocal micrographs of oligodendrocytes (ASPA, white) in the cingulum of the corpus callosum of CD1 mice 7-weeks post-infection. Scale bar 50μm (G and H) Quantification of myelinated axons in the cingulum of the corpus callosum of CD1 mice 7-days post-infection (G) and 7-weeks post-infection (H). n=4 mice per group. (I) Representative transmission electron microscopy (EM) images at the level of the cingulum of the corpus callosum in cross-section. Myelinated axons evident as electron-dense myelin sheaths encircling axons, vewed in cross-section. Scale bars 2μm. Data shown as mean +/- SEM; each dot represents an individual mouse; A,B,D,E,G,H, unpaired, two-tailed t-test. P values shown on figure panels. OPCs: Oligodendrocyte precursor cells. OLGs: Oligodendrocytes.

Consistent with the loss of oligodendrocytes described above, electron microscopy ultrastructural analyses revealed a frank loss of myelin, the insulating ensheathment of axons by oligodendrocyte processes that decreases transverse capacitance and increases the speed of action potential conductance, together with providing metabolic support to axons. We found a decrease in myelinated axon density in subcortical white matter (cingulum of the corpus callosum) evident by 7-days post-infection (Figure 5G-I). This loss of myelin persisted until at least 7 weeks following mild respiratory SARS-CoV-2 infection. Changes in myelin sheath thickness relative to axon diameter (*g*-ratio) were not conclusively identified in the remaining myelin sheaths (Supplementary Figure 6A-D).

Such persistent loss of myelin in subcortical projections would be predicted to impair neural circuit function and axon health, adding to the numerous deleterious neurobiological consequences of SARS-CoV-2 infection.

## Discussion

Taken together, the findings presented here underscore profound multi-cellular dysregulation in the brain caused by even mild respiratory SARS-CoV-2 infection. The white matter-selective microglial reactivity found in both mice and humans following SARS-CoV-2 infection have been shown to inhibit hippocampal neurogenesis, dysregulation of the oligodendroglial lineage and myelin loss in other disease contexts and as demonstrated here. Myelin modulates the speed of neural impulse conduction (Smith and Koles, 1970; Waxman 1980) and provides metabolic support to axons (Funfschilling et al., 2012); even small changes in myelination can exert profound effects on neural circuit dynamics and consequently on cognitive function (Mount and Monje, 2017; Noori et al., 2020; Pajevic et al., 2014). Considering these pathophysiological changes resulting from even relatively mild SARS-CoV-2 infection together with the additional neuropathological consequences that can occur in more severe systemic SARS-CoV-2 infection (Nath and Smith, 2021) - including microvascular thrombi, neuron loss, cortical inflammation, marked and persistent CSF cytokine elevation and even direct brain infection in some cases (Lee et al., 2021; Remsik et al., 2021; Song et al., 2021; Yang et al., 2021) - it is not surprising that the neurological sequalae of COVID-19 infection are proving to be both common and debilitating.

Concordantly, SARS-CoV-2 infection results in a startling rate of persistent symptoms. Long-COVID is thought to affect a high fraction of people who have recovered from SARS-CoV-2 infection. A meta-analysis of 45 studies involving nearly 10,000 subjects found that more than 70% of people experience at least one symptom lasting more than 2 months after infection, with cognitive dysfunction affecting 25% of people, even those who had mild SARS-COV-2 infection (Nasserie et al., 2021). In a study of patients in Italy with COVID-19 pneumonia requiring hospitalization in the spring of 2020, neuropsychometric testing revealed impairment in at least one cognitive domain in 78% of patients at 3-months post-infection, with frequent cognitive impairments affecting psychomotor coordination (57%), executive function (50%), attention and information processing speed (33%), working memory (24%) and verbal memory (10%) (Mazza et al., 2021). Another neuropsychometric study examining all patients with mild, moderate or severe COVID-19 in a New York City hospital system, followed from spring of 2020 through spring of 2021, found impairment in attention (10%), processing speed (18%), memory encoding (24%) and executive function (16%) evident at 7 months after infection. The incidence of cognitive impairment was increased in hospitalized patients compared to those with mild COVID-19; for example, processing speed was impaired in 18% of those with mild COVID-19, and in 28% of those who had COVID-19 severe enough to require hospitalization (Becker et al., 2021). Thus, while those with severe illness are at higher risk, people with even mild COVID-19 may experience lasting cognitive impairment (Tabacof et al., 2020; Tabacof et al., 2022). The incidence and severity of cognitive impairment following COVID-19 caused by newer SARS-CoV-2 variants such as the Omicron variant, or as a result of breakthrough infection in vaccinated individuals, remains to be determined.

Concordant with the clinical similarities in the “COVID-fog” and “chemo-fog” syndromes of cognitive impairment, the findings here illustrate numerous pathophysiological similarities in the multi-cellular dysregulation evident following cancer therapies and following SARS-CoV-2 infection, including white matter microglial reactivity, dysregulation of neural precursor cell populations, reduction in hippocampal neurogenesis, depletion of myelinating oligodendrocytes and myelin loss. As occurs after cancer therapies and in other disease states, the microglial reactivity present after mild respiratory SARS-CoV-2 infection could also induce neurotoxic astrocyte reactivity (Liddelow et al., 2017; Gibson et al., 2019; Guttenplan et al., 2021), as well as aberrant microglial pruning of synapses (Hinkle et al., 2019; Hosseini et al., 2018; Vasek et al., 2016). In the context of severe COVID, microglia in the human cerebral cortex exhibit a transcriptional phenotype that partially overlaps with the neurodegenerative disease-associated microglial phenotype, and cortical astrocytes exhibit a transcriptional program cosnsitent with neurotoxic reactivity (Yang et al., 2021). Whether neurotoxic astrocyte reactivity, induced by the reactive microglia described above (Liddelow et al., 2017; Gibson et al., 2019), occurs after mild respiratory COVID remains to be determined. However, the oligodendrocyte loss observed here could be caused by such neurotoxic astrocyte reactivity, which is known to cause oligodendrocyte cell death (Guttenplan et al., 2021; Liddelow et al., 2017). Microglial depletion strategies can restore similar multi-cellular deficits and rescue impaired cognitive performance in mouse models of cancer therapy-related cognitive impairment (Acharya et al., 2016; Gibson et al., 2019); it remains to be determined if similar therapeutic interventions will prove beneficial for cognitive impairment associated with COVID-19.

Different immune challenges induce diverse immunological responses that differentially involve particular subsets of immune cells and induce diverse cytokine and chemokine profiles systemically and in the CNS. SARS-CoV-2 infection induces a broad inflammatory response – well beyond the typical type 1 immune response seen with other respiratory viral infections (Lucas et al., 2020). Concordantly, we find here that even mild SARS-Cov-2 infection can induce prominent elevations in CCL11, a chemokine known to impair mechanisms of neural plasticity and cognitive function (Villeda et al., 2011), together with white matter-selective microglial reactivity in both mice and humans. In long-COVID patients, most of whom experienced a relatively mild course with their initial SARS-CoV-2 infection, CCL11 was significantly elevated in those with “brain fog” symptoms. In mice, this persistent neuroinflammatory response is associated with impaired hippocampal neurogenesis and loss of both oligodendrocyte precursor cells and mature oligodendrocytes, neuropathological changes predicted by a wealth of previous preclinical work (Geraghty et al., 2019; Gibson et al., 2019; Guttenplan et al., 2021; Liddelow et al., 2017; Monje et al., 2003; Villeda et al., 2011).

The effects of systemic inflammation on microglial reactivity and associated dysregulation of neural cells are not specific to COVID-19. Systemic exposure of rats to low levels of bacterial liposaccharide (LPS) – sufficient to cause mild-moderate sickness behavior but not sepsis - similarly causes microglial reactivity and impairment in hippocampal neurogenesis (Monje et al., 2003). People recovering from critical illness in general experience high rates of long-term cognitive impairment, and persistent cognitive impairment is an important component of post-intensive care syndrome (PICS) (Pandharipande et al., 2013). Other viral syndromes such as influenza cause microglial reactivity and can impair cognitive function, even in the absence of neuroinvasion (Hosseini et al., 2018). However, the particularly immunogenic nature of SARS-CoV-2 (Lucas et al., 2020), together with the magnitude of people infected during this pandemic, contribute to the emerging crisis of persistent cognitive impairment associated with COVID-19 infection. Preclinical studies in disease models that share numerous features with the cellular dysregulation observed here after mild respiratory COVID have identified anti-inflammatory and neuro-regenerative strategies that restore neural plasticity and rescue cognition (Ayoub et al., 2020; Geraghty et al., 2019; Gibson et al., 2019; Monje et al., 2003; Villeda et al., 2011). Such insights from diseases like cancer therapy-related cognitive impairment may elucidate therapeutic strategies to restore healthy cognitive function following COVID-19.

## Acknowledgements

This work was supported by grants from the National Institute of Neurological Disorders and Stroke (R01NS092597 to M.M., NS003130 and NS003157 to A.N.), NIH Director’s Pioneer Award (DP1NS111132 to M.M.), Robert J. Kleberg, Jr. and Helen C. Kleberg Foundation (to M.M.), Cancer Research UK (to M.M.), Waxman Family Research Fund (to M.M. and A.C.G.), National Institute of Allergy and Infectious Diseases (R01AI157488 to A.I.), Fast Grant from Emergent Ventures at the Mercatus Center (to A.I.), RTW Foundation (D.P.) and the Howard Hughes Medical Institute (to A.I. and M.M.).

## STAR Methods

### CONTACT FOR REAGENT AND RESOURCE SHARING

Further information and requests for resources and reagents should be directed to and will be fulfilled by the co-corresponding authors Michelle Monje and Akiko Iwasaki.

## EXPERIMENTAL METHOD AND SUBJECT DETAILS

### Mouse models

6- to 12-week-old female CD1 and BALB/c mice were purchased from Charles River and Jackson Laboratory, respectively, and housed at Yale University. Experiments with wild-type mice transduced with AAV-hACE2 were performed with littermate controls. All procedures used in this study (sex and age matched) complied with federal guidelines and the institutional policies of the Yale School of Medicine Animal Care and Use Committee.

### AAV infection (intratracheal and intracisternal magna injection)

Adeno-associated virus 9 encoding hACE2 (AAV-CMV-hACE2) were made through the HHMI virus core.

### Intratracheal injection

Mice were infected with AAV-hACE2 as previously described (Song et al., 2021). Briefly, animals were anaesthetized using a mixture of ketamine (50 mg/kg) and xylazine (5 mg/kg), injected i.p. The rostral neck was shaved and disinfected. A 5-mm incision was made, the salivary glands were retracted, and the trachea was visualized. Using a 500-μl insulin syringe, a 50-μl bolus injection of 10^11^ genome copies (GC) of AAV-CMV-hACE2 was injected into the trachea. The incision was closed with VetBond skin glue. Following intramuscular administration of analgesic (meloxicam and buprenorphine, 1 mg/kg), animals were placed in a heated cage until full recovery.

### Intracisternal magna injection

Mice were anesthetized using ketamine and xylazine. The dorsal neck was shaved and sterilized. A 2-cm incision was made at the base of the skull, and the dorsal neck muscles were separated using forceps. After visualization of the cisterna magna, a Hamilton syringe with a 15°, 33-gauge needle was used to puncture the dura. 3 μl of AAV_9_ (3 × 10^12^ viral particles/mouse) or mRNA (4– 5 μg) was administered per mouse at a rate of 1 μl/min. Upon completion of the injection, the needle was left in to prevent backflow for an additional 3 min. The skin was stapled and disinfected, and the same postoperative procedures were performed as for intratracheal injections.

### Generation of SARS-CoV-2 stocks

To generate SARS-CoV-2 viral stocks, Huh7.5 cells were inoculated with SARS-CoV-2 isolate USA-WA1/2020 (NR-52281; BEI Resources) to generate a P1 stock. To generate a working VeroE6, cells were infected at an MOI 0.01 for 4 d to generate a working stock. Supernatant was clarified by centrifugation (450 *g* × 5 min) and filtered through a 0.45-μm filter. To concentrate virus, one volume of cold (4°C) 4× PEG-it Virus Precipitation Solution (40% [wt/vol] PEG-8000 and 1.2 M NaCl) was added to three volumes of virus-containing supernatant. The solution was mixed by inverting the tubes several times and then incubated at 4°C overnight. The precipitated virus was harvested by centrifugation at 1,500 *g* for 60 min at 4°C. The pelleted virus was resuspended in PBS and then aliquoted for storage at −80°C. Virus titer was determined by plaque assay using Vero E6 cells.

### SARS-CoV-2 infection

Mice were anesthetized using 30% vol/vol isoflurane diluted in propylene glycol. Using a pipette, 50 μl of SARS-CoV-2 (3 × 10^7^ PFU/ml) was delivered intranasally.

## METHOD DETAILS

### Perfusion and immunohistochemistry

Mice were anesthetized with isoflurane and transcardially perfused with 10ml 0.1M PBS followed by 10ml 4% paraformaldehyde (PFA). Brains were fixed in 4% PFA overnight at 4°C, before being transferred to 30% sucrose for cryoprotection. Brains were embedded in Tissue-Tek (Sakura) and sectioned coronally at 40μm using a sliding microtome (Microm HM450; Thermo Scientific). For immunohistochemistry, a 1 in 6 series of 40μm coronal sections was incubated in 3% normal donkey serum in 0.3% Triton X-100 in TBS blocking solution at room temperature for one hour. Rabbit anti-SARS-CoV-2 nucleocapsid (1:250, GeneTex GTX135357), rabbit anti-IBA1 (1:1000, Wako Chemicals 019-19741), rat anti-CD68 (1:200, Abcam ab53444), rabbit anti-DCX (1:500, Abcam ab18723), rabbit anti-ASPA (1:250, EMD Millipore ABN1698), goat anti-PDGFRa (1:500; R&D Systems AF1062), mouse anti-CC1 (1:100; EMD Millipore OP80) were diluted in 1% normal donkey serum in 0.3% Triton X-100 in TBS and incubated overnight at 4°C (CC1 was incubated for 7 days at 4°C). Sections were then rinsed three times in 1X TBS and incubated in secondary antibody solution (Alexa 647 donkey anti-rabbit IgG, 1:500 (Jackson Immunoresearch); Alexa 488 donkey anti rat IgG, 1:500 (Jackson Immunoresearch); Alexa 594 donkey anti-rabbit IgG, 1:500 (Jackson Immunoresearch); Alexa 488 donkey anti goat IgG, 1:500 (Jackson Immunoresearch); Alexa 647 donkey anti-mouse IgG, 1:500 (Jackson Immunoresearch) in 1% blocking solution at room temperature protected from light for two hours. Sections were rinsed 3 times in TBS, incubated with DAPI for 5 minutes (1:1000; Thermo Fisher Scientific) and mounted with ProLong Gold mounting medium (Life Technologies).

### Confocal imaging and quantification

All cell counting was performed by experimenters blinded to experimental conditions on a Zeiss LSM700 or LSM800 scanning confocal microscope (Zeiss). Images were taken at 20X magnification and analyzed using FIJI software. For microglia imaging, three consecutive sections ranging approximately from bregma +1.2 to +0.8 were selected for cortex and corpus callosum, and four consecutive sections ranging approximately from bregma -1.8 to -2.4 were selected for the hilus of the dentate gyrus. For each section, the superficial cortex, deep cortex, cingulum, genu of the corpus callosum, and hilus of the dentate gyrus were identified, and two 219.5 × 219.5 um fields per slice were selected in those areas for quantification. All CD68^+^ and IBA1^+^ cells were counted in those regions. For analysis of immature neurons, four consecutive sections ranging approximately from bregma -1.8 to -2.4 were selected for the dentate gyrus, and four 219.5 × 219.5 μm fields per slice were selected in those areas for quantification. All DCX^+^ cells within the inner layer of the dentate gyrus were counted in those regions. For analysis of oligodendrocytes (ASPA^+^ or CC1^+^) and oligodendrocyte precursor cells (PDGFRa^+^) four to five consecutive sections ranging from bregma +1.2 to +0.8 were stained. The cingulum of the corpus callosum in each hemisphere was imaged with one 319.5 × 319.5 μm field and cells were quantified using the “Spots” function in Imaris image analysis software (Oxford Instruments). Each image was manually inspected to ensure accurate oligodendrocyte counts. False positive and false negative counts were manually removed or added, respectively.

### Electron Microscopy

7 days or 7 weeks after infection (as described above), mice were sacrificed by transcardial perfusion with Karnovsky’s fixative: 2% glutaraldehyde (Electron Microscopy Sciences (EMS 16000) and 4% paraformaldehyde (EMS 15700) in 0.1 M sodium cacodylate (EMS 12300), pH 7. Samples were prepared as detailed previously in (Gibson et al., 2014). Axons in the cingulum of the corpus callsoum were analyzed for *g*-ratios calculated by dividing the shortest axonal diameter by the total diameter (diameter of axon/diameter of axon plus myelin sheath), as well as for myelinated axon density (total number of myelinated axons in each frame) by experimenters who were blinded to experimental conditions and genotypes. A minimum of 300 axons were scored per group.

### Mouse CSF and serum cytokine analysis

The Luminex multiplex assay was performed by the Human Immune Monitoring Center at Stanford University. Mouse 48-plex Procarta kits were purchased from Thermo Fisher, Santa Clara, California, USA, and used according to the manufacturer’s recommendations with modifications as described below. Briefly: beads were added to a 96-well plate and washed in a Biotek ELx405 washer. Samples were added to the plate containing the mixed antibody-linked beads and incubated overnight at 4°C with shaking. Cold (4°C) and room temperature incubation steps were performed on an orbital shaker at 500-600 RPM. Following the overnight incubation plates were washed in a Biotek ELx405 washer and then biotinylated detection antibody added for 60 minutes at room temperature with shaking. Plate was washed as described above and streptavidin-PE was added. After incubation for 30 minutes at room temperature, washing was performed as above and reading buffer was added to the wells. CSF samples were measured as singlets, while serum samples were measured in duplicate. Plates were read on a FM3D FlexMap instrument with a lower bound of 50 beads per sample per cytokine. Custom Assay Chex control beads were purchased from Radix Biosolutions, Georgetown, Texas, and were added to all wells.

### Isolation of patient plasma and cytokine/chemokine measurements

Patient plasma was collected with informed consent and IRB approval as previously described (Lucas et al., 2020). Patient whole blood was collected in sodium heparin-coated vacutainers. Blood samples were gently agitating at room temperature until delivered to Yale from Mount Sinai Client. All blood samples were processed on the day of collection. Plasma samples were collected following centrifugation of whole blood at 400*g* without brake at room temperature for 10 minutes. The undiluted plasma was then transferred to 15 ml polypropylene conical tubes, aliquoted, and stored at −80 °C. Plasma samples were shipped to Eve Technologies (Calgary, Alberta, Canada) on dry ice, and levels of cytokines and chemokines were measured using the Human Cytokine Array/Chemokine Array 71-403 Plex Panel (HD71). All samples were measured at the time of the first thaw.

### Human immunohistochemistry

Human tissue samples were prepared and analyzed as previously described (Lee et al., 2021). Human brains obtained at the time of autopsy in New York and in Iowa during the spring of 2020 were fixed in formalin. Five-micron thickness sections were obtained from formalin-fixed paraffin-embedded (FFPE) blocks. Slides were deparaffinized and then rehydrated using xylene and graded ethanol. Preparation of slides for immunohistochemistry involved antigen retrieval by Heat-Induced Antigen Retrieval (HIAR) methods in 10 mM citrate buffer at pH 6.0 or 10 mM Tris/EDTA buffer at pH 9.0, or Enzyme-Induced Antigen Retrieval (EIAR) methods in proteinase K. Peroxidase blocking was done by incubating tissues in 0.3% to 3.0% hydrogen peroxide for 10 minutes at room temperature. Protein blocking was performed using protein block solution (DAKO) for 30 minutes at room temperature. Sections were exposed to primary antibodies and incubated overnight at room temperature. Antibodies included IBA1 (Wako 019-19741) and CD68 (Thermo Fisher Scientific 14-0688-82). Following 1x tris-buffered saline (TBS) wash containing 0.05% TritonX-100, sections were incubated with PowerVision polymeric horseradish peroxidase (poly-HRP) anti-mouse IgG (Leica Biosystems) or PowerVision polyHRP anti-rabbit IgG for 2 hours at room temperature. Antibody binding was visualized with 3,3′- diaminobenzidine (DAB; Vector Laboratories). Sections were then counterstained with hematoxylin (DAKO). Slides were coverslipped using EcoMount mounting medium (Biocare Medical). A whole slide scanner (Aperio AT2, Leica Biosystems) was used for image acquisition.

### QUANTIFICATION AND STATISTICAL ANALYSIS

For all quantifications, experimenters were blinded to sample identity and condition.

For fluorescent immunohistochemistry and confocal microscopy, images were taken at 200X and cells considered co-labeled when markers co-localized within the same plane. For cortical/white matter microglia staining three sections per mouse, eight frames in standardized locations per section (24 images per mouse) were counted, for hippocampal microglia staining, four sections per mouse, 2 frames in standardized location per section (8 images per mouse) were counted. For hippocampal neuroblast staining, four sections per mouse, 4 frames in standardized location per section (16 images per mouse) were counted. For oligodendrocyte staining four to five sections per mouse, 2 frames in standardized location per section (8-10 images per mouse) were mounted. For each mouse, 300-900 cells were counted for each immunohistochemical marker analysis. The density of cells was determined by dividing the total number of cells quantified for each lineage by the total volume of the imaged frames (mm^3^). In all experiments, “n” refers to the number of mice. In every experiment, “n” equals at least 3 mice per group; the exact “n” for each experiment is defined in the figure legends.

For electron microscopy (EM) quantification of *g*-ratio and myelinated axon density, 6000X images were obtained. *g*-ratios were measured by dividing the axonal diameter by the diameter of the entire fiber (diameter of axon/diameter of axon + myelin sheath) using ImageJ software. For myelinated axon density, the number of axons surrounded by a myelin sheath were counted and normalized to tissue area. 29 images were scored for each mouse. Approximately 400-700 axons were scored for each mouse.

All statistics were performed using Prism Software (Graphpad). For all analyses involving comparisons of one or two variables, 1-way or 2-way ANOVAs, respectively, were used with Tukey’s multiple comparisons post hoc corrections used to assess main group differences. For analyses involving only two groups, unpaired two-tailed Student’s t tests were used. Shapiro-Wilk test was used to determine normality for all datasets; all datasets were parametric. A level of p < 0.05 was used to designate significant differences.

---

**Supplementary Figure 1.**
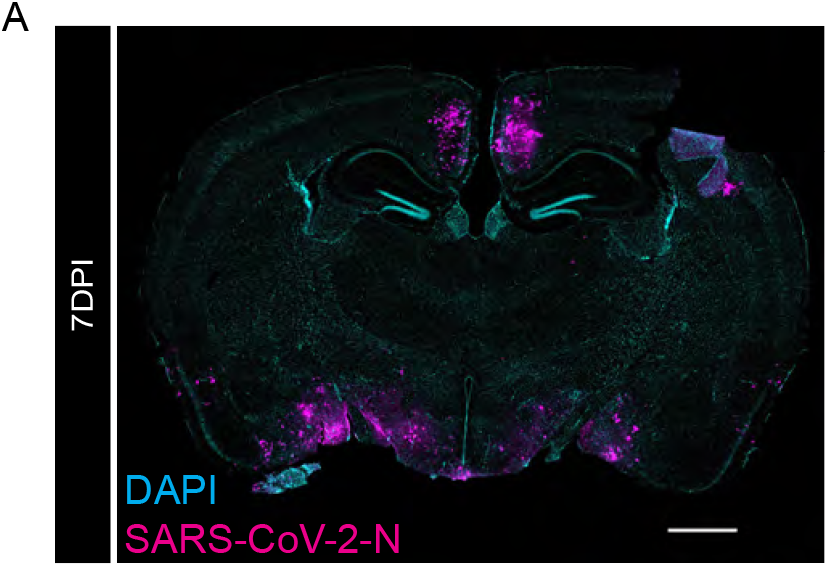
Neuroinvasive SARS-CoV-2 mouse model as a positive control for nucleocapsid protein immunostaining. (A) Confocal micrograph of coronal section of mouse brain, illustrating SARS-CoV-2 nucleocapsid protein (SARS-CoV-2-N) 7-days post-infection (SARS-CoV-2-N, magenta; DAPI, cyan). Scale bar 1mm. Related to Figure 1.

**Supplementary Figure 2.**
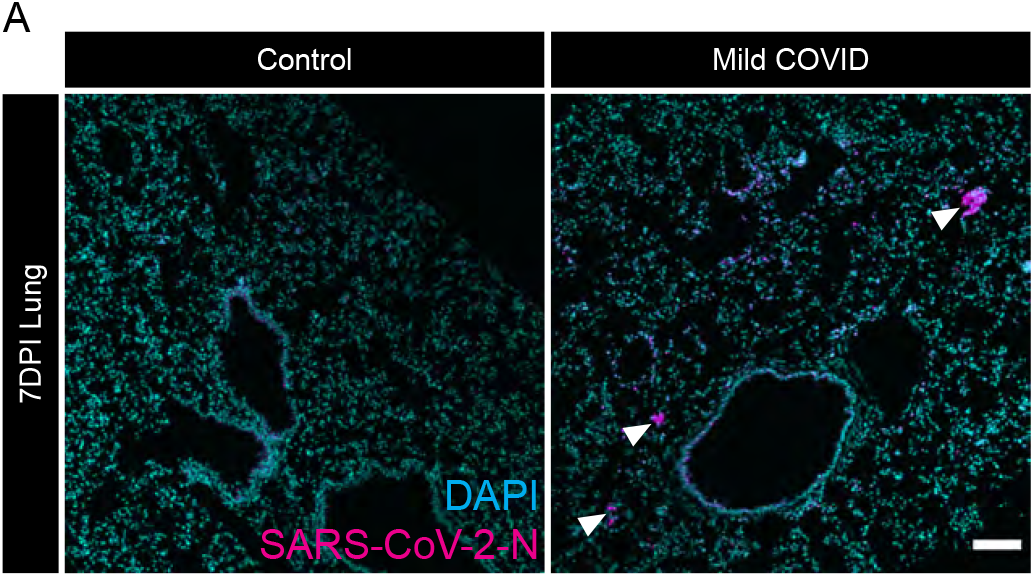
Evidence of SARS-CoV-2 infection in lung of mild respiratory COVID mouse model. (A) Representative confocal micrographs of SARS-CoV-2 nucleocapsid protein (SARS-CoV-2-N, magenta; DAPI, cyan) in mouse lung 7-days post-infection. Arrowheads highlight SARS-CoV-2-N nucleocapsid protein immunostaining. Scale bar 100μm. Related to Figure 1.

**Supplementary Figure 3.**
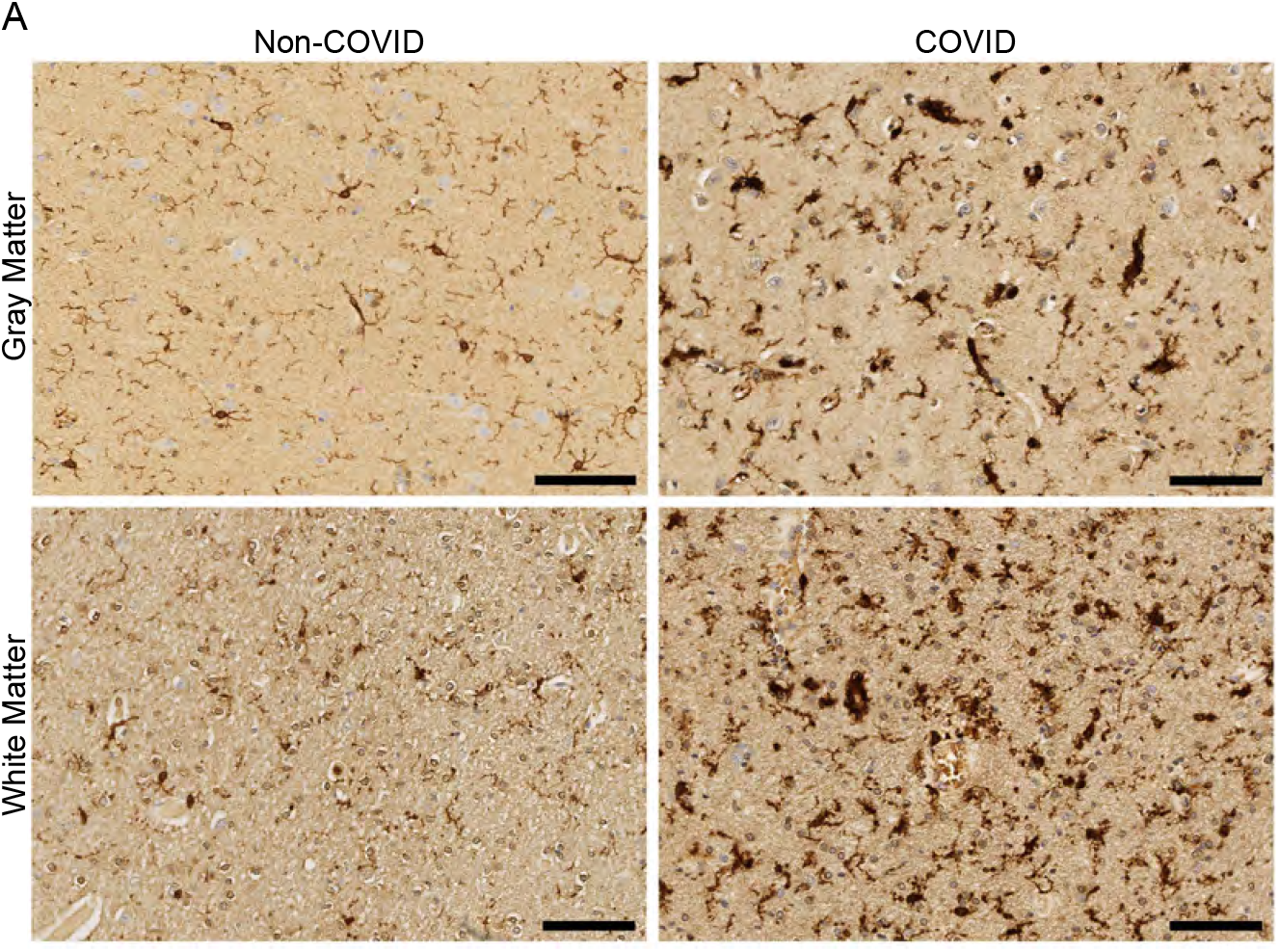
White matter-selective microglial reactivity in humans with SARS-CoV-2 infection. (A) Representative micrographs of IBA1 immunostaining (brown) in the cerebral cortex (gray matter) or subcortical white matter of human subjects with or without COVID. Scale bars 100μm. Related to Figure 3.

**Supplementary Figure 4.**
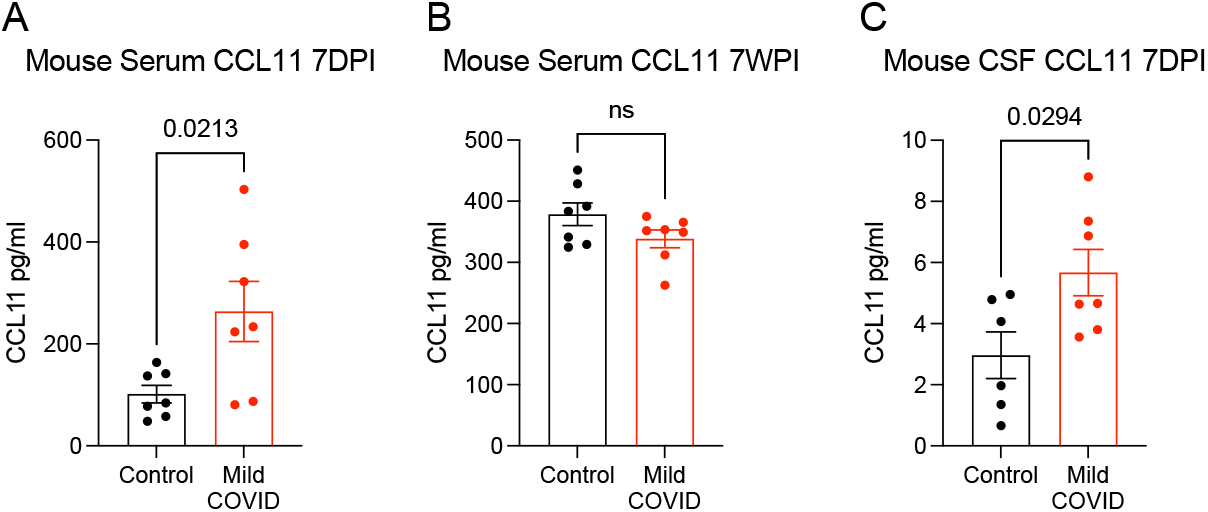
CCL11 levels after mild respiratory SARS-CoV-2 infection. (A and B) Serum levels of CCL11 from CD1 mice 7-days post-infection (A) and 7-weeks post-infection (B). n=7 mice per group. (C) CCL11 levels in CSF of CD1 mice 7-days post-infection. n=7 mice per group. Data shown as mean +/- SEM; each dot represents an individual mouse; P values shown on figure panels; ns p>0.05; two-tailed unpaired t-test. Data shown in heatmap form in Figure 1. Related to Figures 1 and 4.

**Supplementary Figure 5.**
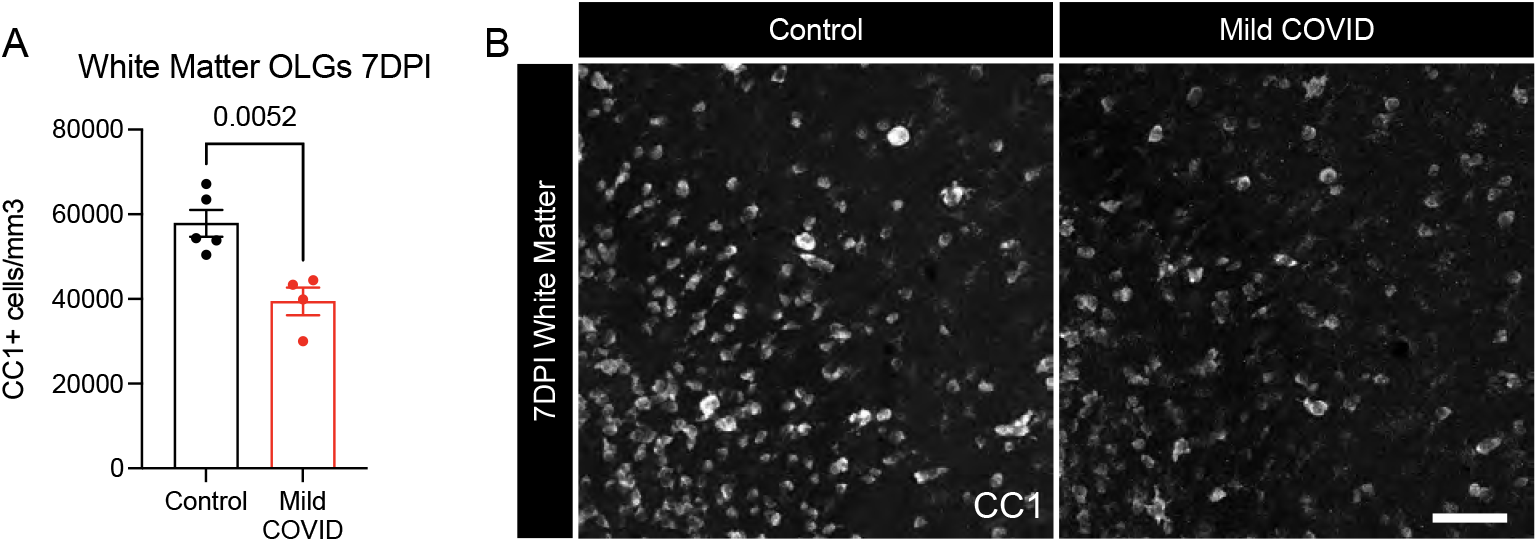
Validation of oligodendrocyte loss after mild respiratory SARS-CoV-2 infection. (A) Oligodendrocyte (CC1+) quantification in the cingulum of the corpus callosum of CD1 mice 7-days post-infection. (B) Representative confocal micrographs of oligodendrocytes (CC1, white) in the cingulum of the corpus callosum of CD1 mice 7-days post-infection. Scale bar 50μm. Data shown as mean +/- SEM; each dot represents an individual mouse; unpaired, two-tailed t-test. P value shown on figure panel. Related to Figure 5.

**Supplementary Figure 6.**
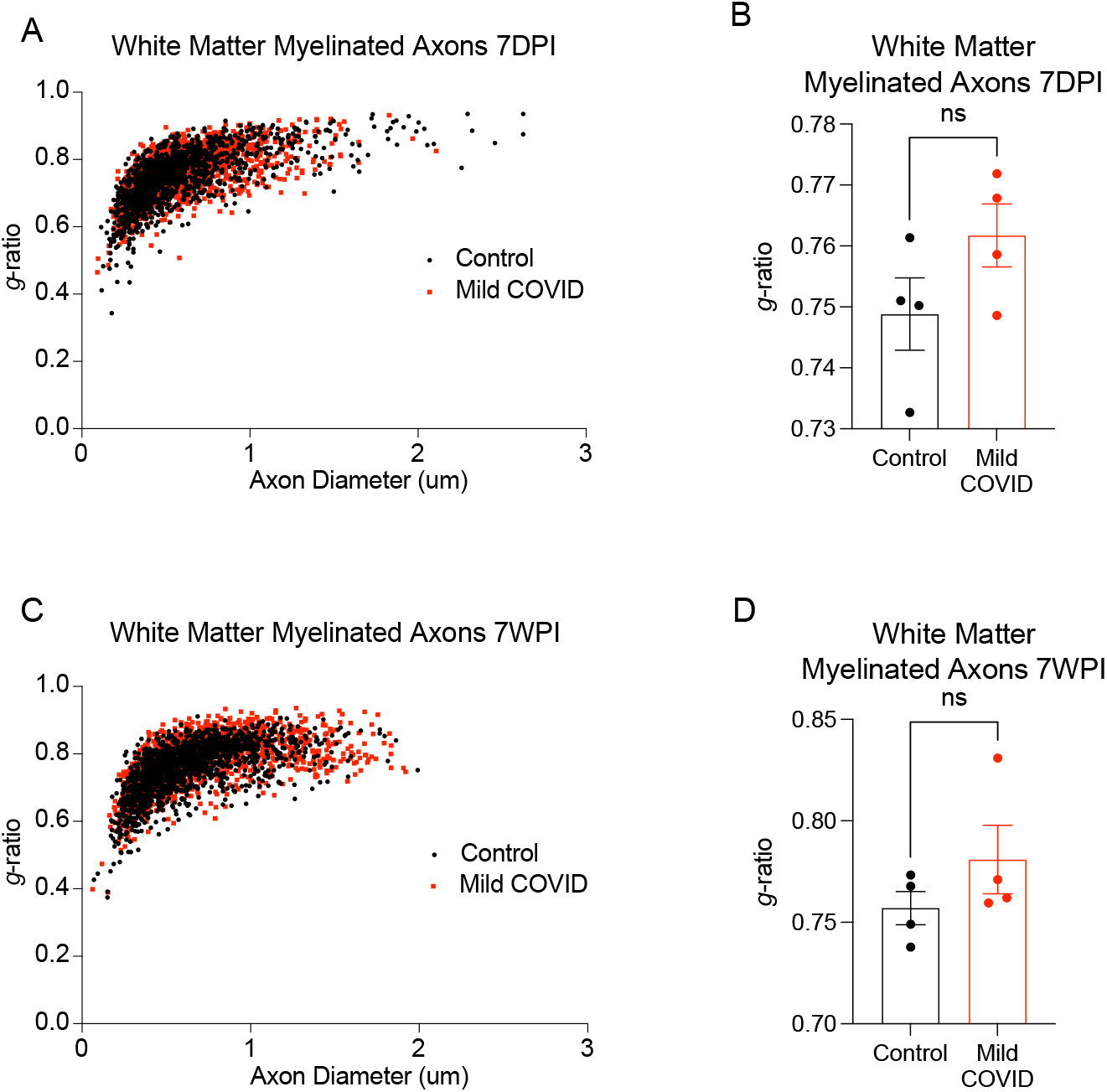
Myelin sheath thickness after mild respiratory SARS-CoV-2 infection. (A) Scatter plots of *g*-ratio relative to axon diameter 7-days-post infection. Black dots, control axons; red dots, axons from mice with mild respiratory COVID. (B) Cumulative *g*-ratios of myelinated axons per animal at 7-days-post infection. Each dot represents an individual mouse. (C) Scatter plots of *g*-ratio relative to axon diameter 7-weeks-post infection. Black dots, control axons; red dots, axons from mice with mild respiratory COVID. (D) Cumulative *g*-ratios of myelinated axons per animal at 7-weeks-post infection. Each dot represents an individual mouse. Data in B and D shown as mean +/- SEM; n=4 mice per group; ns p>0.05 by two-tailed, unpaired t-test. Related to Figure 5.

**Supplementary Table 1.**
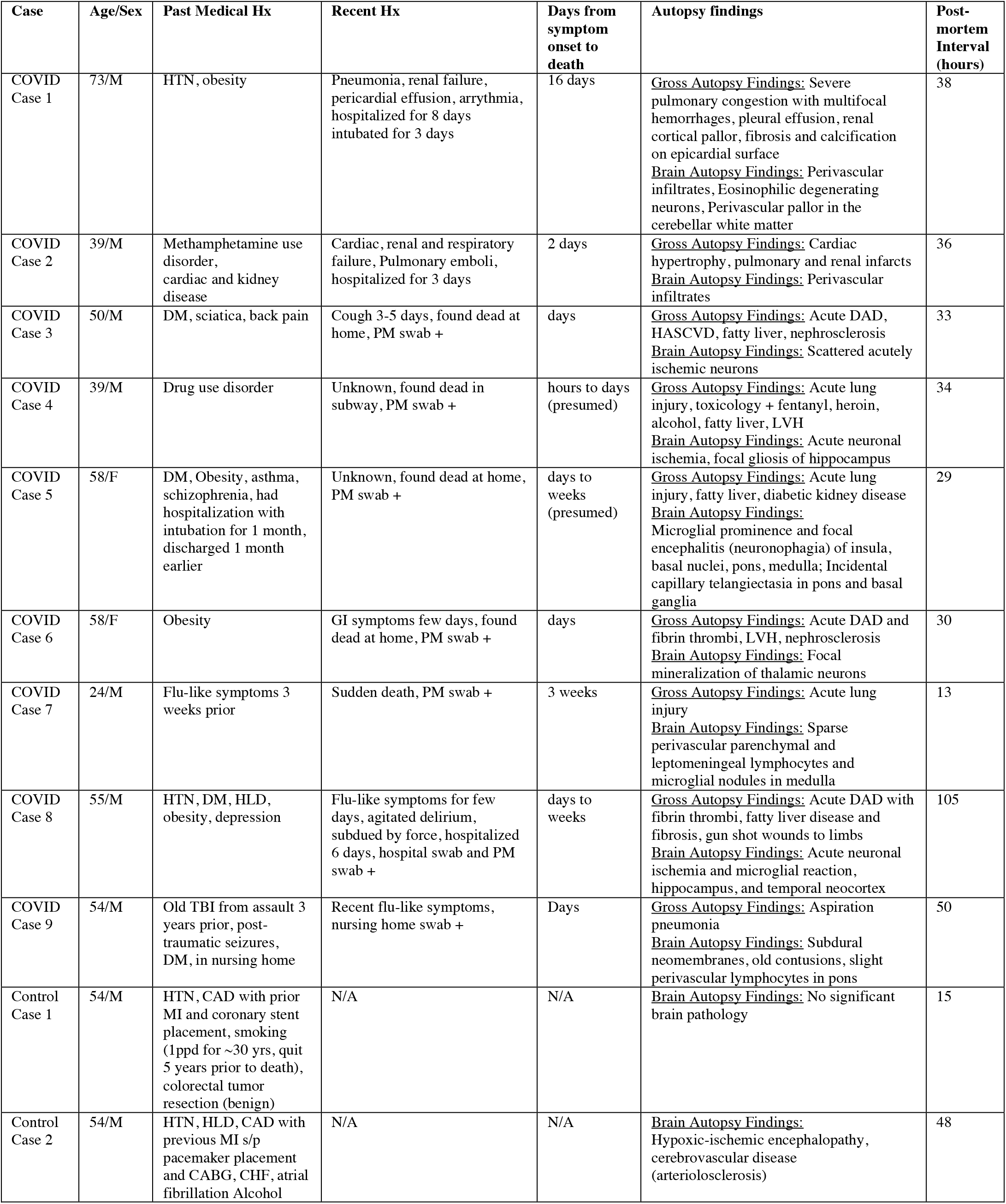
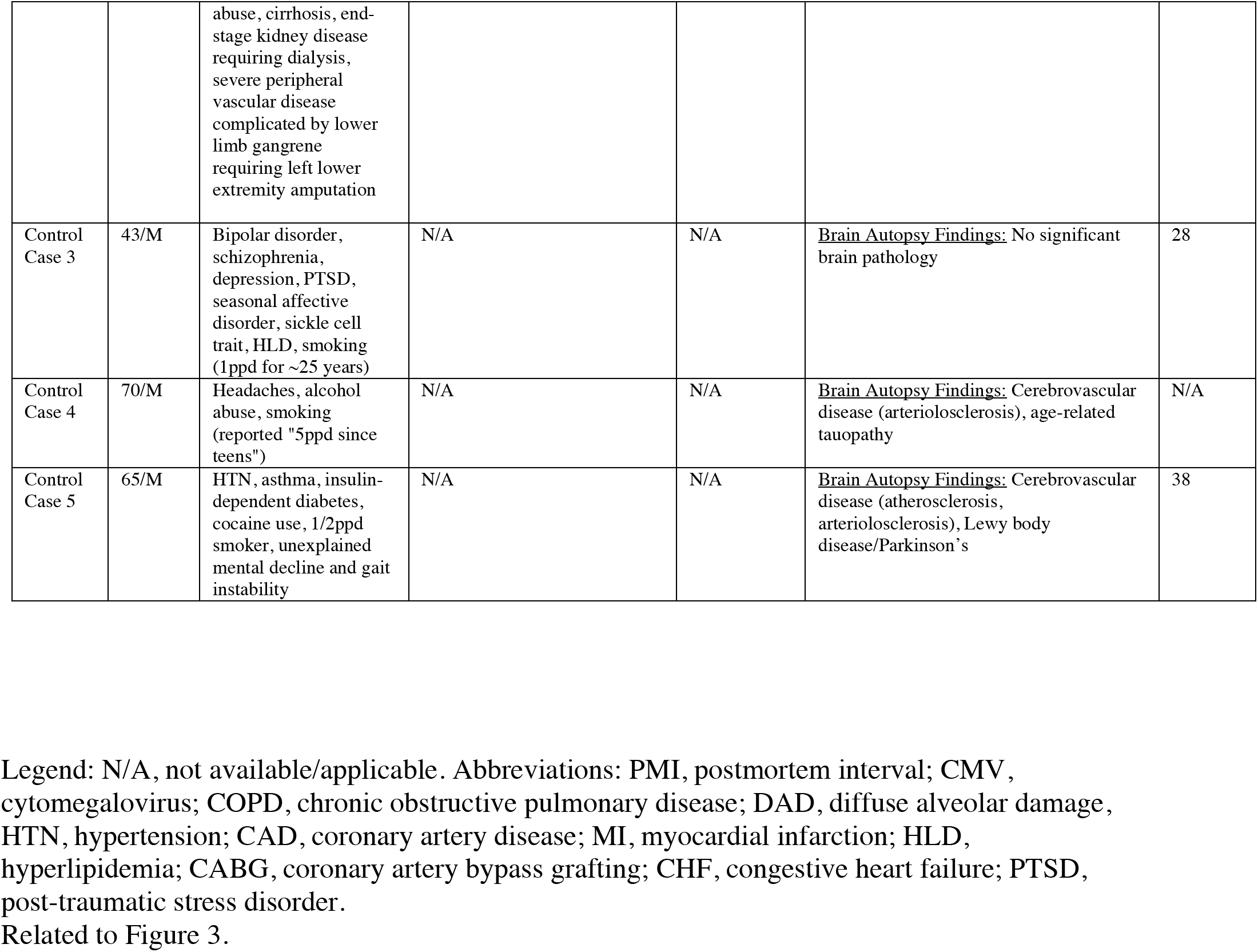
COVID-19 subject and non-COVID-19 control subject characteristics. Related to Figure 3.

